# Eoulsan 2: an efficient workflow manager for reproducible bulk, long-read and single-cell transcriptomics analyses

**DOI:** 10.1101/2021.10.13.464219

**Authors:** Nathalie Lehmann, Sandrine Perrin, Claire Wallon, Xavier Bauquet, Vivien Deshaies, Cyril Firmo, Runxin Du, Charlotte Berthelier, Céline Hernandez, Cédric Michaud, Denis Thieffry, Stéphane Le Crom, Morgane Thomas-Chollier, Laurent Jourdren

**Author notes:** Co-first authors (Nathalie Lehmann, Sandrine Perrin).

## Abstract

**Motivation:** Core sequencing facilities produce huge amounts of sequencing data that need to be analysed with automated workflows to ensure reproducibility and traceability. Eoulsan is a versatile open-source workflow engine meeting the needs of core facilities, by automating the analysis of a large number of samples. Its core design separates the description of the workflow from the actual commands to be run. This originality simplifies its usage as the user does not need to handle code, while ensuring reproducibility. Eoulsan was initially developed for bulk RNA-seq data, but the transcriptomics applications have recently widened with the advent of long-read sequencing and single-cell technologies, calling for the development of new workflows.

**Result:** We present Eoulsan 2, a major update that (i) enhances the workflow manager itself, (ii) facilitates the development of new modules, and (iii) expands its applications to long reads RNA-seq (Oxford Nanopore Technologies) and scRNA-seq (Smart-seq2 and 10x Genomics). The workflow manager has been rewritten, with support for execution on a larger choice of computational infrastructure (workstations, Hadoop clusters, and various job schedulers for cluster usage). Eoulsan now facilitates the development of new modules, by reusing wrappers developed for the Galaxy platform, with support for container images (Docker or Singularity) packaging tools to execute. Finally, Eoulsan natively integrates novel modules for bulk RNA-seq, as well as others specifically designed for processing long read RNA-seq and scRNA-seq. Eoulsan 2 is distributed with ready-to-use workflows and companion tutorials.

**Availability and implementation:** Eoulsan is implemented in Java, supported on Linux systems and distributed under the LGPL and CeCILL-C licenses at: http://outils.genomique.biologie.ens.fr/eoulsan/. The source code and sample workflows are available on GitHub: https://github.com/GenomicParisCentre/eoulsan. A GitHub repository for modules using the Galaxy tool XML syntax is further provided at: https://github.com/GenomicParisCentre/eoulsan-tools

## Background

For the last fifteen years, technological advances in sequencing devices have resulted in a dramatic increase in read throughput. Furthermore, the rise of long-read sequencing with the third generation sequencers from Pacific Biosciences (PacBio) and Oxford Nanopore Technologies (ONT) enables the sequencing of much longer fragments. Altogether, the panel of High Throughput Sequencing (HTS) applications is very large, with a current democratization of single-cell approaches.

A common issue with HTS data, is to ensure reproducibility and traceability of the bioinformatics analyses performed on sequencing results, regardless of the application (research, medical diagnostics or forensics). Analysis pipelines encapsulate a series of sequential steps, for which a variety of individual tools must be connected. Such pipelines can be written as simple scripts, or with more elaborate workflow management systems. Sequencing core facilities are particularly in need of reliable automated pipelines that do not require manual intervention, while ensuring proper traceability.

Since its inception, the aim of Eoulsan (Jourdren et al. 2012) is to provide an efficient, stable and reliable workflow engine supporting HTS analyses, targeting bioinformatician end-users, and especially core facilities that analyze several dozen of projects each year. To do so, Eoulsan’s core design encodes the analysis workflow within an XML flat file, rather than embedding it within the code of a program. Another core feature is the proper separation between the experimental design (samples and associated metadata) and the analysis workflow itself. This model allows to reuse and standardize workflows across many experiments. Moreover, Eoulsan avoids file manipulation issues, by using file aliases linked to external repositories (e.g., for genomes, features annotations or mapper genome indexes). It automatically checks the input data on startup, and prevents output files overriding. Eoulsan is distributed with state-of-the-art individual tools (e.g., FASTQC, MultiQC, DESeq2) embedded in reusable modules, and offers alternative choices for many of the steps (e.g., mapping can be performed with either BWA, Bowtie1, Bowtie2, STAR, GMAP, GSNAP or MiniMap2).

Eoulsan supports diverse computing infrastructures to parallelize and distribute computation - from workstations to large clusters - thereby ensuring its efficiency and versatility. Our software was one of the very first bioinformatic tools able to run on an Hadoop cluster. The aim of Hadoop is to apply algorithms where data is physically stored, rather than sending data to computing nodes hosting the algorithms. In genomics, sequencing data are often stored in quite huge files. On a typical scientific cluster, transferring data to the computing nodes is thus problematic, with a risk of staturating access to network file systems, resulting in using a lower number of nodes. A Hadoop cluster solves this input/output (I/O) issue, enabling the use of the maximum available number of nodes. Hadoop thus remains particularly suited for genomics applications.

Several workflow engines have been developed specifically for genomics (recently reviewed in (Wratten, Wilm, and Göke 2021)). Among them, the most widely used are Galaxy (Afgan et al. 2018), Snakemake (Mölder et al. 2021) and Nextflow (Di Tommaso et al. 2017). Eoulsan’s main principle is its low-code approach, which is closer to Galaxy’s design. Its strength also lies in its installation ease, the stability of the code, and continuous maintenance and evolution for the last 10 years.

Here, we present Eoulsan 2, a major update of our workflow engine software with many new features and enhancements. This new version facilitates the development of new modules, includes module containerization, and extends its execution environments with support for several job schedulers. Novel modules have also been added to use state-of-the-art third-party tools. Eoulsan 2 supports transcriptomics applications, including both short and long reads bulk differential analyses, and single-cell RNA-seq workflows for Smart-seq2 and 10x Genomics. Moreover, we provide several ready-to-use workflows and tutorials for common analyses to help users to start with our workflow engine, accessible from the GitHub page of the project [https://github.com/GenomicParisCentre/eoulsan].

## Implementation

Eoulsan is a free software published under the GNU LGPL and CeCCIL-C licenses. A Java Runtime Environment under Linux is the only requirement of this tool. Its unique user interface is the command line. Eoulsan can be run under three modes: local (workstation), Hadoop cluster or “standard” cluster (SLURM, TORQUE, HT-Condor and PBSPro job schedulers are supported). In cluster mode, users can define memory and processor count requirements for each task. Moreover, merger and splitter steps can be added to the workflow to better scale data processing all over the cluster and speedup computation.

Only two text files describing the pipeline need to be provided by the user to the input of the workflow engine: (i) the experimental design and (ii) the workflow definition (Figure 1). The experimental design of an Eoulsan analysis is stored in a text file (see example in Figure 2). For Eoulsan 2, the design file format has been enhanced to handle complex designs for DESeq2 differential gene analysis (Love, Huber, and Anders 2014), such as multiple comparisons and DESeq2 design formulas. The workflow steps and theirs parameters are listed in a companion XML file, separated from the code, ensuring flexibility and traceability (see example on the GitHub project page: https://raw.githubusercontent.com/wiki/GenomicParisCentre/eoulsan/files/workflow-rnaseq.xml). Altogether, these two files allow to quickly resume large analyses upon trouble-shooting, and guarantees reproducibility.

**Figure 1:**
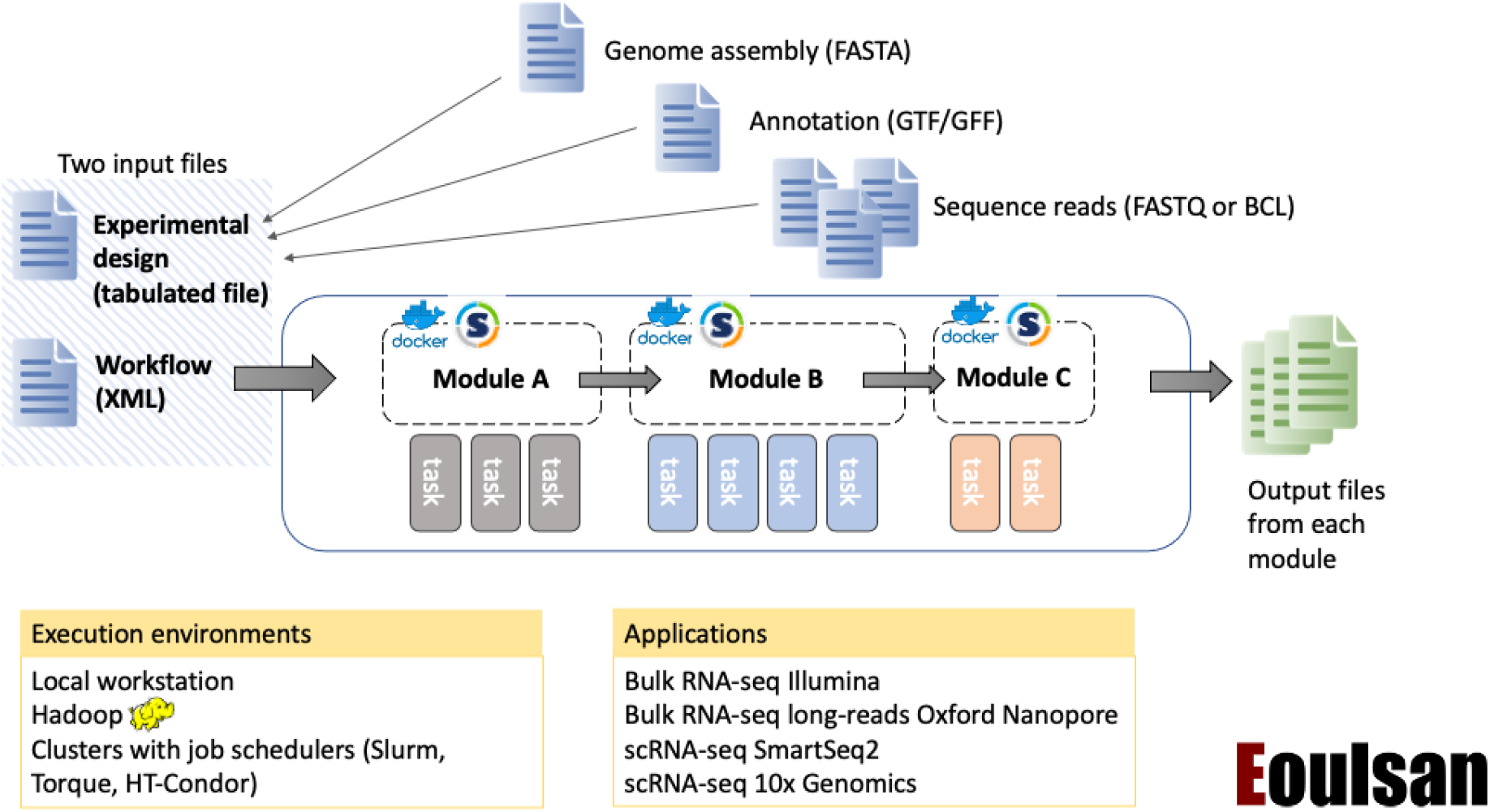
Overview of Eoulsan 2 workflow management system. Eoulsan is a workflow engine designed for high-throughput genomics analyses, with a focus on transcriptomics. The workflow is described as an XML file, rather than code, and each step corresponds to a reusable independent module. Each module may internally run several tasks with third-party programs. Installation of these dependencies is facilitated by containerization with Docker or Singularity. Implementation of a new module is facilitated by the use of Galaxy Tools XML (not shown). The experimental design file specifies the metadata of the samples, as well as their status (controls), and links to larger files that may be stored elsewhere in the filesystem, such as the genome assembly and annotation. The workflow can be executed either on a local workstation or on a cluster, which enables parallelization of the tasks when possible. Ready-to-use workflows are available for four applications.

**Figure 2:**
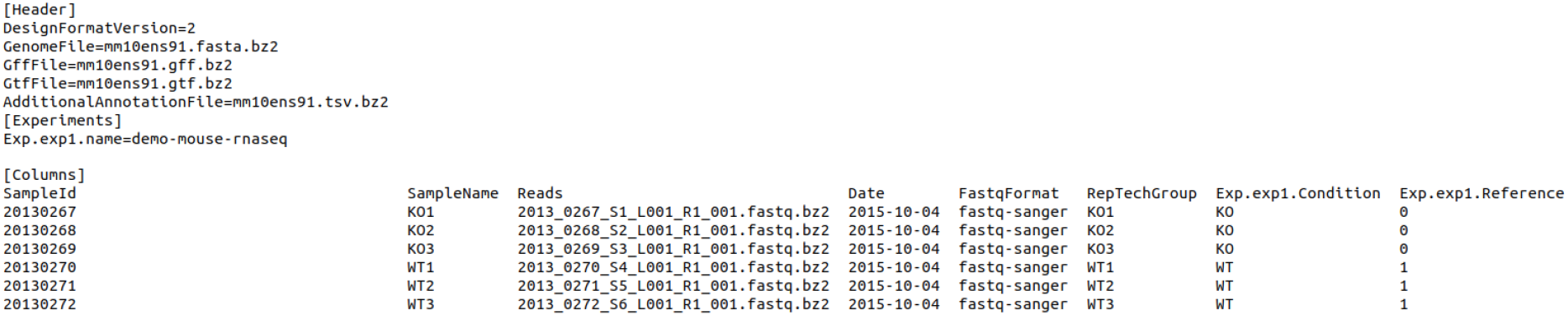
Example of a design file for Eoulsan. The header section contains general information about the design and the project. This includes the genome file that will serve for the mapping, and the annotation. The “Experiments” section specifies details on the experimental design. It can include multiple experiments and comparisons that should be made for the differential analyses. The “Columns” section provides details on each sample, including the information about technical and biological replicates. The file can be found on the GitHub project for Eoulsan at this URL: https://raw.githubusercontent.com/wiki/GenomicParisCentre/eoulsan/files/design-rnaseq.txt

In the first Eoulsan release, the bundled RNA-seq workflow was hardcoded in the software, thus not allowing to easily add new steps. In Eoulsan2, we have rewritten the workflow engine, making it versatile for any bioinformatics workflow to be implemented.

In Eoulsan, each step of the workflow is a module, corresponding to encapsulated third-party tools (Figure 1). In addition to previous modules, Eoulsan2 is shipped with established tools such as DESeq2 (Love, Huber, and Anders 2014), HTSeq-count (Anders, Pyl, and Huber 2015), sam2bam [https://github.com/samtools/htsjdk], FastQC [https://www.bioinformatics.babraham.ac.uk/projects/fastqc/], MultiQC (Ewels et al. 2016), Trimmomatic (Bolger, Lohse, and Usadel 2014)…). External modules can be added through Java plugins. In an effort to simplify the development of a new module, Eoulsan 2 is now able to handle modules defined with the Galaxy tool XML syntax. Numerous modules have been adapted using this syntax (e.g., Cutadapt (Martin 2011), featureCounts (Liao, Smyth, and Shi 2014), UMI-Tools (Smith, Heger, and Sudbery 2017), …), and are easily accessible on a dedicated GitHub repository [https://github.com/GenomicParisCentre/eoulsan-toolsl. It is thus easy to adapt an existing program from Galaxy to Eoulsan.

To ensure reproducibility of results over multiple computers, Docker [https://www.docker.com/] has streamlined container management. Eoulsan can rely on Docker or Singularity [https://sylabs.io/singularity/] containers for both Galaxy Tool XML and Java modules (e.g., DESeq2). Each module thus runs separately in its own container. This approach simplifies the deployment of the modules and avoids dependency management and complex installation procedures.

Since the first version of our tool, the number of available modules has dramatically increased (10 modules initially vs 33 modules, among which 14 are built with the Galaxy tool syntax and available on our specific GitHub repository). To ensure consistency of the code over versions, we have deployed a functional test system based on Jenkins [https://www.jenkins.io/] continuous integration software. This system checks every week whether changes in Eoulsan source code has led to result changes in more than 190 reference analyses (2 days of analyses). The tests cover all modules and Galaxy tools, mapping programs, as well as local and Hadoop execution modes.

With unit tests for Java source code, functional tests and container management, Eoulsan 2 ensures a very strong confidence in reproducibility of user results.

## RESULTS

While the first version of Eoulsan was designed only for bulk RNA-seq differential analysis workflows, Eoulsan 2 is shipped with ready-to-use workflows for various types of transcriptomics analyses:

- bulk RNA-seq Illumina differential analysis
- Oxford Nanopore long reads RNA-seq
- scRNA-seq with Smart-seq2 protocol
- scRNA-seq with 10x Genomics protocol

Ready-to-use workflows, along with tutorials detailing their usage, are accessible on the GitHub page of Eoulsan [https://github.com/GenomicParisCentre/eoulsan].

### Bulk RNA-seq Illumina differential analysis

In its first version, Eoulsan supported four read mappers (BWA, Bowtie, SOAP2, GSNAP). In Eoulsan 2, the RNA-seq workflow has been greatly improved with new integrated mappers (STAR, Bowtie2, GMAP/GSNAP), while SOAP2 is no longer supported. We have developed a fast implementation of the HTSeq-count algorithm in Java, which is included in Eoulsan 2. It also supports complex designs with DESeq2.

### Bulk Oxford Nanopore long reads RNA-seq differential analysis

The Oxford Nanopore long-reads RNA-seq workflow uses Minimap2 (Li 2018) to align sequences over the reference genome. The workflow is otherwise very similar to the Illumina workflow, with the differential analysis performed with DESeq2. In all the steps (except for mapping), the workflow uses the same modules as the Illumina workflow, with a tuning of the parameters to adapt to the long-reads. The steps with tuned parameters are: filter raw FASTQ files (filterreads module), remove unmap and multimatches SAM entries (filtersam module) and count alignments with HTSeq-count (expression module). Even though the workflow currently relies on the same tools as used for the Illumina workflow, we are expecting to update the workflow once the LRGASP challenge reaches its conclusions [https://www.gencodegenes.org/pages/LRGASP/]. This systematic evaluation of different methods for transcript computational identification and quantification will constitute a reference to help us identify the most relevant tools, and integrate them as new modules in Eoulsan.

### Single-cell RNA-seq pre-processing analyses

To answer user growing interest for single-cell transcriptomics data, we developed scRNA-seq workflows for both Smart-seq2 and 10x Genomics technologies. These workflows focus on the pre-processing steps, from the raw reads up to the generation of the count matrix. These workflows are detailed in Figure 3. These first steps are crucial to ensure the quality of the pre-processed data before the downstream analyses.

**Figure 3:**
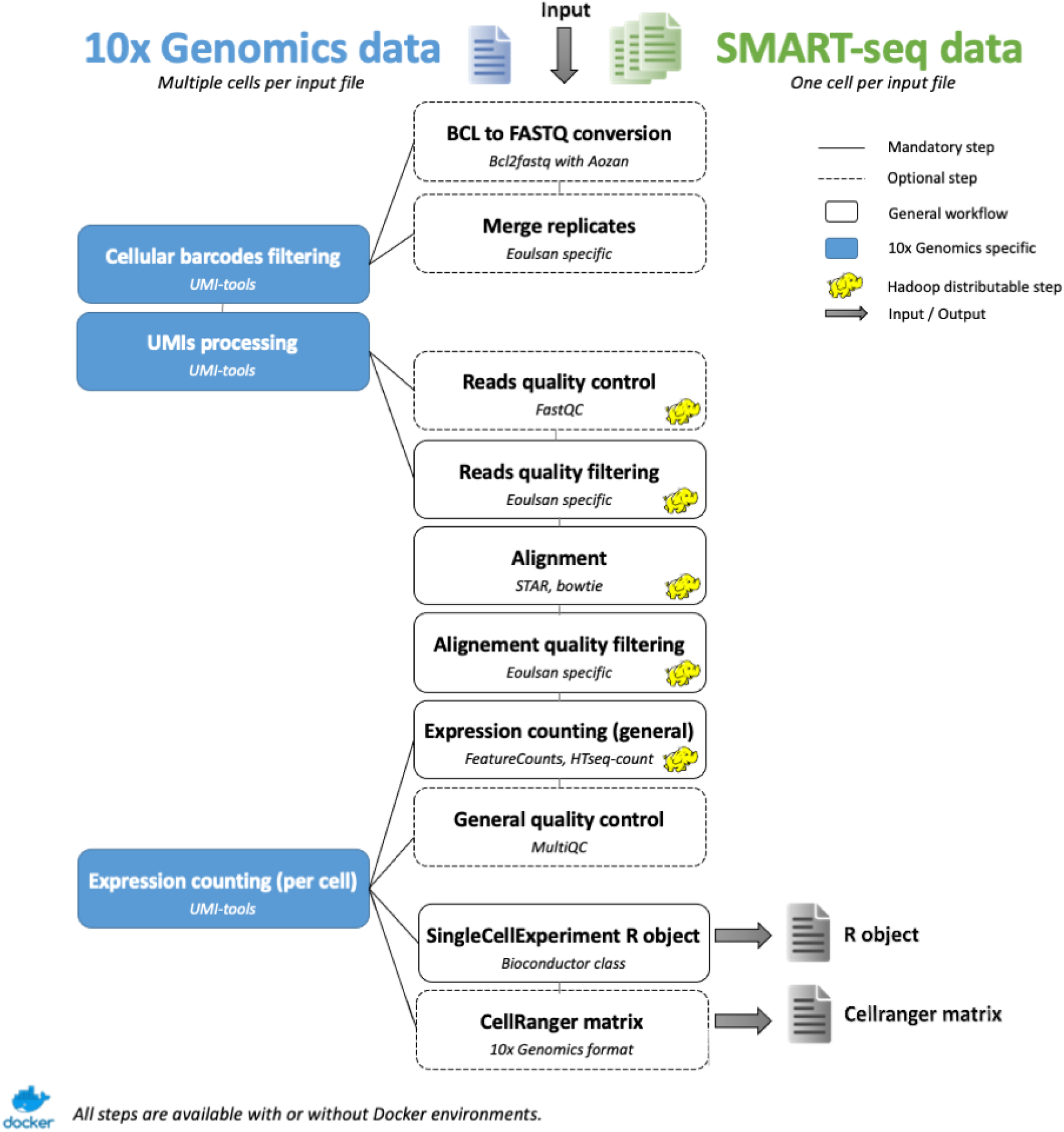
Single-cell RNA-seq workflows in Eoulsan 2. Eoulsan 2 supports two workflows for scRNA-seq data processing: Smart-seq2 and 10x Genomics. Some modules were specifically developed to support this type of data. While the Smart-seq2 workflow is quite similar to a bulk RNA-seq workflow, the 10x Genomics workflow has several specific modules to treat cell barcodes and Unique Molecular Identifiers (UMIs).

The Smart-seq2 workflow is derived from the bulk RNA-seq Illumina workflow, as the steps are quite similar, with the mapping performed by STAR. The main difference is that data for each cell is stored in a separate FASTQ file. A typical experiment thus produces hundreds of FASTQ files. The fact that Eoulsan uses Hadoop calculation cluster makes this tool well-suited for parallelized processes, hence handling this issue.

The 10x Genomics workflow reuses some existing modules of Eoulsan (e.g., mapping with STAR), but has required the development of novel modules to take into account the cellular barcodes and Unique Molecular Identifiers (UMIs), intrinsic to this technology. The 10x Genomics protocol uses cell barcodes (16bp) to identify each cell. Many barcodes do not correspond to real cells, which means “valid” barcodes corresponding to real cells must be inferred. For these steps, we rely on the third-party tool UMI-Tools (Smith, Heger, and Sudbery 2017). By default, UMI-tools whitelist detects these valid barcodes with the knee method, by retaining the top most abundant barcodes (umiwhitelist module).

Following the identification of these « valid » barcodes, the next step is to filter out the reads that do not match with these barcodes (umiextract module). Once the mapping has been performed, reads must be assigned to the genes they most probably originated from. This step is handled with the featureCounts module (Liao, Smyth, and Shi 2014), which outputs a bulk count matrix. Finally, the single-cells counts are isolated with the umicount module, thus producing a single-cell expression matrix. It should be noted that only the reads mapping to the same strand as the annotated genes are taken into account, since the 10x Genomics protocol is stranded.

Downstream analyses are generally directly performed in R. Eoulsan facilitates this transition to post-processing steps with R, by generating a RDS file containing a SingleCellExperiment Bioconductor object. It encapsulates the count matrix, along with cells and gene annotations, as a ready-to-use object for downstream analyses. Another possibility is to output a CellRanger formatted count matrix with the MatrixToCellRangerMatrix module. This option allows users to perform downstream analyses in R with functions that take as input a CellRanger matrix. Altogether, Eoulsan 2 ensures users don’t have to change their downstream analyses pipeline without further changes, by directly providing the count matrix in the correct format.

### Comparison with similar tools

In terms of Workflow Management Systems, Nextflow and Snakemake are currently the most common engines for a usage at the command line, and Galaxy for a usage through a web browser interface. Eoulsan’s originality lies in its internal design as a low-code workflow manager. Indeed, the workflows use XML files without any code. The modules are actually an abstraction layer over the actual tools, thereby hiding the complete syntax of all their parameters. This organisation enables to maintain the code of the modules separately from the workflow itself. This choice is particularly suited in the context of a core facility platform that analyses dozens of projects each year. Each project is thus associated with a reproducible workflow file, independent of the command lines that are actually run. Portability of the code is comparable to the above-mentioned workflow engines, as many modules are containerized with Docker or Singularity, and we take advantage of systems developed by Galaxy (Galaxy Tools XML syntax). In addition, the code consistency in Eoulsan is tested with functional tests using a Jenkins server on a weekly basis.

Our 10x Genomics scRNA-seq workflow can be compared to the CellRanger program, developed by the same company. CellRanger performs the pre-processing steps, and provides a user-ended HTML report already including some downstream analyses (clustering, t-SNE visualisation). The Quality Check (QC) values are limited in CellRanger, while Eoulsan provides FASTQC and MultiQC reports that enables a deeper interpretation of the quality of the data. Eoulsan relies on state-of-the-art tools such as UMI-Tools, which enables more versatility than CellRanger. CellRanger’s lack of time efficiency and high requirement for memory usage has previously been underlined (Gao et al. 2021). Our workflow is 2-3 times faster with limited resource usage for similar results (R2=0.998 for UMI count per cell) (Figure 4).

**Figure 4:**
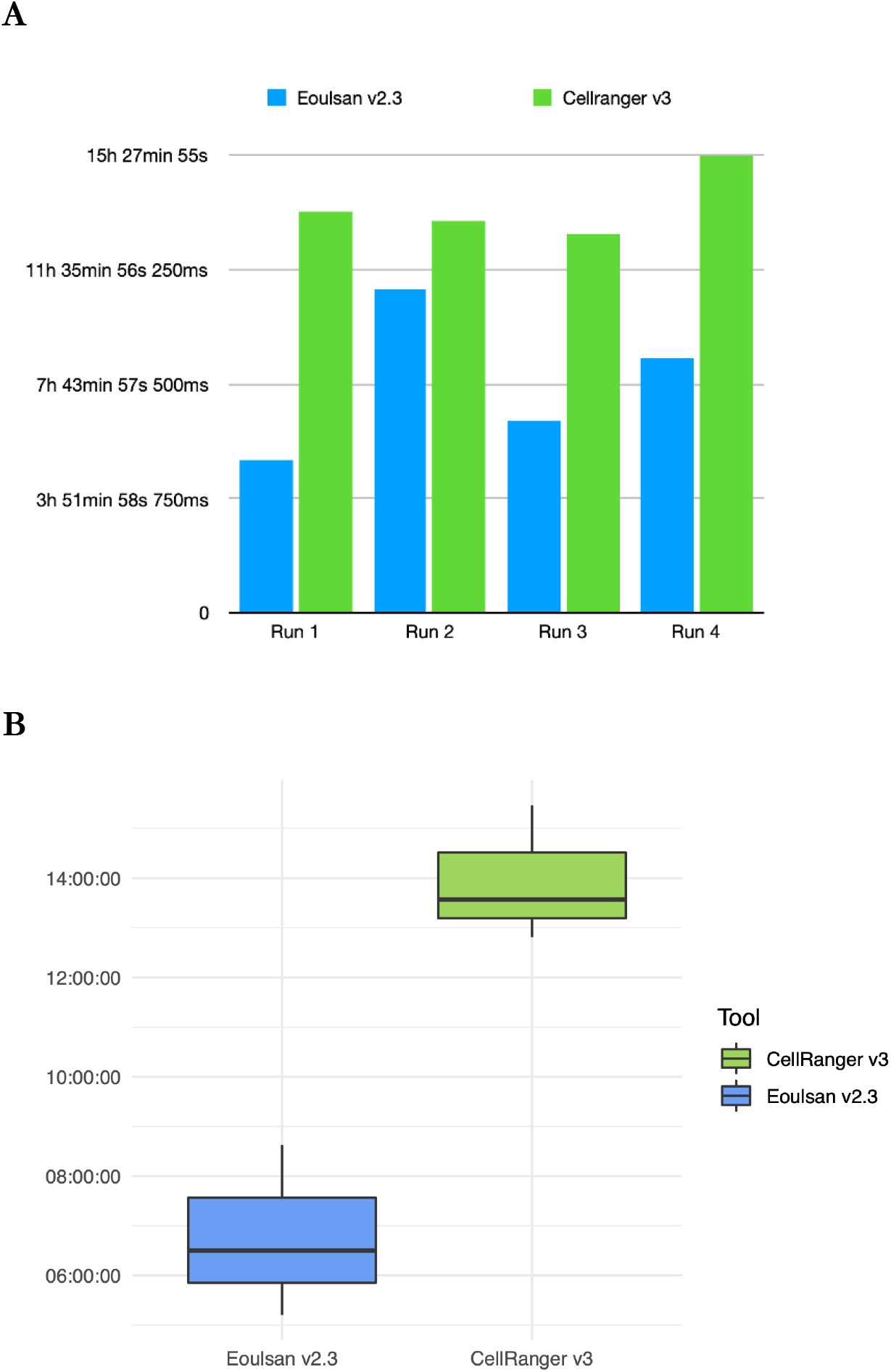
Comparison of computing efficiency between Eoulsan and CellRanger. The comparison was performed using Eoulsan v2.3 and CellRanger v3, on a dataset of 5000 PBMC cells generated with the 10x Genomics technology (v3 Chemistry) downloaded from 10x Genomics website: [https://www.10xgenomics.com/resources/datasets/5-k-peripheral-blood-mononuclear-cells-pbm-cs-from-a-healthy-donor-v-3-chemistry-3-1-standard-3-0-2]. **A**. Barplot of each of the 4 individual runs. For comparison purposes with CellRanger that uses a pre-computed index, we ran Eoulsan with a pre-computed STAR index, except for run 2. Note that for run 2, Eoulsan is faster than CellRanger, even when adding the step of building the STAR index. **B**. Boxplot showing the execution time (Y axis) for each program, summarizing the three runs using a pre-computed index.

## Conclusions

Our workflow engine Eoulsan 2 aims at facilitating high throughput sequencing analysis for bioinformaticians, in particular for transcriptomics applications. Eoulsan handles the most resource-expensive parts of analyses, and it can conveniently be deployed on any cluster, as well as on workstations. Result reproducibility has always been one of our major concerns in Eoulsan development. This is why Eoulsan source code is extensively tested through unit tests, as well as a huge number of functional tests. In addition, since Docker and Singularity container systems have emerged, they have been utilized for Eoulsan modules external dependencies. To ease module creation, we introduced in Eoulsan 2 the support for the Galaxy Tool XML files, and a comprehensive documentation for developers. Moreover, several ready-to-use workflows for both short and long reads RNA-seq, Smart-seq2 and 10x Genomics scRNA-seq are available, along with tutorials. With all these enhancements since its first version ten years ago, and a strong foundation for reproducibility and scalability, Eoulsan is particularly suited for core facilities wishing to implement a long-term stable solution for managing their transcriptomics workflows.

## Availability and requirements

### Project name

Eoulsan

### Project home page

https://www.outils.genomique.biologie.ens.fr/eoulsan/: Binary downloads and reference documentation

https://github.com/GenomicParisCentre/eoulsan: Source code and workflow wiki

https://github.com/GenomicParisCentre/eoulsan-tools: Additionnal modules

### Operating system(s)

Unix

### Programming language

Java

### License

GNU LGPL and CeCCIL-C licenses

### Any restrictions to use by non-academics

none

## Acknowledgements

We thank Hugo Varet for helping with DESeq2 usage, Sophie Lemoine for intensive application testing and feedbacks, Pierre-Marie Chiaroni for the initial work on modules of the ChIP-seq pipeline, Aurelien Birer for help with two Galaxy tools, Geoffray Brelurut for helping with the Smart-seq pipeline, Hatim El Jazouli for testing the 10x single-cell pipeline, and Bpipe authors for the job scheduler submission scripts. We also thank the IBENS informatics and bioinformatics core facilities for helping with testing Eoulsan on the cluster, as well as the TGCC platform for testing on their cluster.

## Contributions

N.L developed, tested, documented and compared the 10x Genomics workflow. S. P developed the Galaxy tools integration and functional test system. C.W ported HTSeq-count to Java and Hadoop. X.B implemented the new design file format and support for DESeq2. V.D rewritten DESeq 1 support. C.F created sam2fastq, bam2sam modules and enhanced XLSX/ODS export system. R.D integrated many read filters using Trimmomatic. C.B improved design file checks for DESeq2. C.H supervised C.M who developed modules for format interconversion bed/bigbed/bam/bigwig and the trimadapt module. D.T. was involved in student supervision and provided ideas for the general development of Eoulsan. S.L.C. and M.T.C. were involved in proposing and planning the addition of new features, and were involved in the supervision of students and engineers. L.J. developed the first version of Eoulsan, rewrote the workflow engine, supervised and coordinated the addition of new modules and features, and ensured the long-term maintenance of the project L.J. and M.T.C. drafted the manuscript, with input from N.L, S.L.C and D.T.

